# hafeZ: Active prophage identification through read mapping

**DOI:** 10.1101/2021.07.21.453177

**Authors:** Christopher J. R. Turkington, Neda Nezam Abadi, Robert A. Edwards, Juris A. Grasis

## Abstract

**Summary:** Bacteriophages that have integrated their genomes into bacterial chromosomes, termed prophages, are widespread across bacteria. Prophages are key components of bacterial genomes, with their integration often contributing novel, beneficial, characteristics to the infected host. Likewise, their induction—through the production and release of progeny virions into the surrounding environment—can have considerable ramifications on bacterial communities. Yet, not all prophages can excise following integration, due to genetic degradation by their host bacterium. Here, we present hafeZ, a tool able to identify ‘active’ prophages (i.e. those undergoing induction) within bacterial genomes through genomic read mapping. We demonstrate its use by applying hafeZ to publicly available sequencing data from bacterial genomes known to contain active prophages and show that hafeZ can accurately identify their presence and location in the host chromosomes.

**Availability and Implementation:** hafeZ is implemented in Python 3.7 and freely available under an open-source GPL-3.0 license from https://github.com/Chrisjrt/hafeZ. Bugs and issues may be reported by submitting them via the hafeZ github issues page.

## 1 Introduction

Bacteriophages, viruses that infect bacteria, are heavily involved in many aspects of bacterial physiology and evolution. Temperate bacteriophages in particular, are tightly interwoven into bacterial biology through their ability to integrate their DNA into their host’s chromosome via the lysogenic cycle of bacteriophage replication (Howard-Varona et al., 2017). When integrated, bacteriophages are called prophages and can contribute fitness changes to their hosts through expanding the hosts’ genetic repertoire (e.g. by providing toxin or antibiotic resistance genes); or by altering the expression of existing bacterial genes (e.g. by integrating directly into a bacterial gene and causing its disruption) (Ofir and Sorek, 2018).

However, prophages not only influence bacterial biology when integrated into the genome, they also influence bacterial behaviour through their induction. Here, prophages excise from the chromosome and begin virion production via the lytic cycle of bacteriophage replication—a process that is often fatal for the host bacterium. Through induction to the lytic cycle, temperate bacteriophages have been associated with major changes in the densities of both lysogenised and non-lysogenised bacterial populations, alterations in bacterial biofilm formation levels, and changes in bacteria-host interactions in host-associated systems (Keen and Dantas 2018).

Given their importance, numerous computational tools have been developed in recent years to identify integrated bacteriophages within bacterial genomes. To date though, most prophage identification tools are unable to determine whether an identified prophage can be induced and no tool utilises prophage induction as a parameter for their detection. This is particularly important because prophages are subject to decay in their host (usually by the accumulation of transposases), wherein some prophages will lose their ability to excise and produce viral particles, leaving them as dormant regions of the bacterial genome (Bobay et al. 2014). Here we present hafeZ a tool that identifies active prophages (i.e. those undergoing induction) in bacterial genomes by examining the mapping of genome sequencing data to an assembly for signs of prophage induction. The workflow of hafeZ is described below and a summarised overview can be found in figure S1.

## 2 Description

### 2.1 Inputs

hafeZ requires three main inputs from the user: (1) a complete/contiguous genome assembly in FASTA format; (2) a set of reads for the given genome in FASTQ format; and (3) the path to the folder containing the Prokaryotic Virus Orthologous Groups (pVOGs; Grazziotin et al., 2017) database that is downloaded during the initial setup of hafeZ.

### 2.2 Read mapping and region of interest identification

hafeZ begins by examining the number and size of contigs in an assembly and removing small contigs (default = < 10,000 bp). Contigs passing this filter are then indexed and the reads are mapped to the contigs using minimap2 (Li, 2018), with mapping results converted to coverage depths using samtools (Li et al., 2009) and mosdepth (Pedersen and Quinlan, 2018). Coverage values are then smoothed using a Savitzky-Golay filter via the ‘*savgol_filter*’ function of SciPy (Virtanen et al., 2020) to reduce instances of short lapses in coverage depth in otherwise heavily covered regions.

Once coverage values have been collected and smoothed, the modified Z-score of coverage per base across the length of the genome are then calculated using the equation of Iglewicz and Hoaglin, 1993:

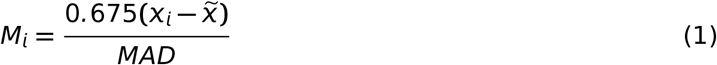

Where *M_i_* is the modified Z-score of a base’s coverage, *x_i_* is the coverage of that base, 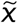 is the median of all coverage values, and *MAD* is the median absolute deviation. *MAD* is the median of absolute deviations about the median for all per-base coverage values and is calculated as:

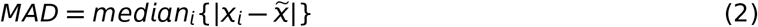

The modified Z-score values are then used to identify regions within the genome with higher than expected coverage via numpy (Harris et al., 2020). Regions are called if they pass thresholds for minimum modified Z-score (default = 3.5) and region width (default = 4,000 bp). Regions passing these filters within a close vicinity of each other are then merged to create a ‘region of interest’ (ROI; figure S2).

To identify a potential deletion event, the reads mapping within each ROI, and those mapping within 15,000 bp either side of it, are extracted using pysam (https://github.com/pysam-developers/pysam). For ROIs located centrally in a contig (defined as any ROI > 15,000 bp from either end of a contig) reads are examined for the presence of at least one read pair where read partners map a distance of roughly the length of the ROI from each other. This would indicate that their sequenced fragment includes a case where the prophage has excised, as the read pair exists closer than would be expected for the assembly (figure S3-A). If no distant reads are found, the ROI is dropped from further analysis. For ROIs passing this threshold and ROIs located near the end of a contig, the reads are then searched for any soft clipped reads (reads where only part of a read maps to a given location; figure S3-B). For each ROI, assembly positions where at least 10 reads have been soft clipped are collected. Then, all combinations of these locations that occur on opposite sides of the ROI center are collated. The collated combinations then serve as the basis for putative prophage start and end locations for each ROI to then be further filtered.

### 2.3 Region of interest filtering

Each combination of start-end locations are then filtered by examining the modified Z-scores in and around those locations. If a prophage induction event had occurred, it would be expected that the region between the start and end positions would have a median modified Z-score value greater than the Z-score threshold, while in the regions preceding the start and proceeding the end locations the median modified Z-scores should be below the Z-score threshold as these should be bacterial chromosome regions. Therefore, hafeZ first removes any ROI start and end location combinations where this is not the case. For ROIs located at the start/end of a contig, only the region between the ROI start/end and the sides not near contig ends are used in filtering. The best combinations of each soft clipping location passing the Z-score filter are retained (default = best 50), using the sum of start/end soft clipped read count as the scoring metric (higher = better).

As plasmids would pass all filters to this point, hafeZ then examines any ROI start/end combinations near the ends of a contig for indications that the contig is circular (i.e. reads mapping from the end of a contig to the start and vice versa). If true, these ROIs are then flagged as circular but are carried forward as they may also be extra-chromosomal prophages.

To further-filter ROI start/end combinations, hafeZ then maps the clipped portion of soft clipped reads for all non-circular ROIs and examines their mapping location. Here, hafeZ examines if the clipped portion of the read maps near the position of where the reads on the opposing side of the ROI were clipped (figure S3-C). This step adds additional stringency to hafeZ but can be disabled by using the ‘*-N/--no_extra*’ option.

The single best start/end combination for each ROI is then collected, with the best start/end combination being that with the highest sum of soft clipped reads at the start/end plus the number of clipped sections from these mapping near the opposing position.

### 2.4 Sequence analysis

After ROI coordinates have been determined, genes are then predicted using Pyrodigal (https://github.com/althonos/pyrodigal), a cythonised version of Prodigal (Hyatt et al., 2010), and the sequences of the predicted open reading frames (ORFs) are then extracted. ROIs containing less than a minimum number of ORFs (default = 6) are then removed, with the peptide sequences encoded by each gene of the surviving ROIs then compared to the pVOGs database using hmmscan (HMMER v3.3.1; http://hmmer.org). ROIs are retained if the proportion of ORFs in the ROI showing similarity to a pVOGs hidden Markov model (HMM) profile passes a user-defined threshold (default = 0.1 i.e. 10% of ORFs). The *att* sites for each of ROI are then determined by extracting the region ± 100 bp either side of the ROI start/end locations and examining them for homology using BLASTn (Camacho et al. 2009) with the settings ‘*-evalue 10000 -task blastn-short*’. The putative *att* sites with the lowest e-values and a length > 11 bp are then output as the potential *att* sites for each ROI.

### 2.5 Output

hafeZ generates all outputs in the path provided by the user. If an ROI passes all filters, six main outputs are produced: (1) a multi-FASTA file containing the DNA sequence of each ROI identified; (2) a fasta file for each ROI containing the DNA sequences of all ORFs; (3) a fasta file for each ROI containing the amino acid sequences of all ORFs; (4) a tab-separated file containing details of ORFs hit by the pVOGs comparisons; (5) a tab-separated summary file containing key information on all identified putative prophage regions; and (6) a figure showing Z-score distributions for each contig found to contain a ROI with the location of the ROI highlighted (e.g. figure S4). If no ROIs are found, or no ROIs pass the filtering process, only an empty tab-separated summary file will be output.

### 2.6 Example usage

In their work, Zünd et al., 2021 used a combination of comparing the coverage mapping between induced vs non-induced samples and read examination on 14 bacterial isolates to identify 10 active prophages. To illustrate the ability of hafeZ to identify active prophages from bacterial sequencing data, we applied hafeZ to this sequencing data-set (European Nucleotide Archive project No. PRJEB39818) using default settings and the corresponding publicly available reference assemblies as mapping targets for each read-set (table S1). We found that overall hafeZ was able to identify all 10 prophages in the data-set with no deviation in predicted start/end locations compared to those of Zünd et al., 2021. However, as hafeZ analyses individual samples for the presence of prophages and the Zünd data-set contains triplicate induced and non-induced samples that were compared to identify their prophages in their work, the presence or absence of each of these prophages was dependant on the sample. hafeZ did identify one element not mentioned by Zünd, corresponding to a plasmid in *Salmonella enterica* serovar Typhimurium LT2^*p*22^. This plasmid was flagged as circular in all cases though, and thus can be easily identified as such. A table summarising the presence/absence and positional differences for each expected prophage can be found in table S2, while the combined outputs of hafeZ for all samples can be found in table S3.

## 3 Conclusion

We show that hafeZ is highly accurate at detecting active prophages in bacterial genomes and that it is able to identify prophages in a diverse range of samples and organisms. We believe hafeZ is an ideal tool to be used in addition to current integration-focused prophage identification tools to give highly accurate prophage positions, describe the activity of prophages identified by other tools, and, through its novel use of induction as an identification metric, could potentially identify novel viruses that would be missed by existing prophage detection algorithms.

---

**Figure S1:**
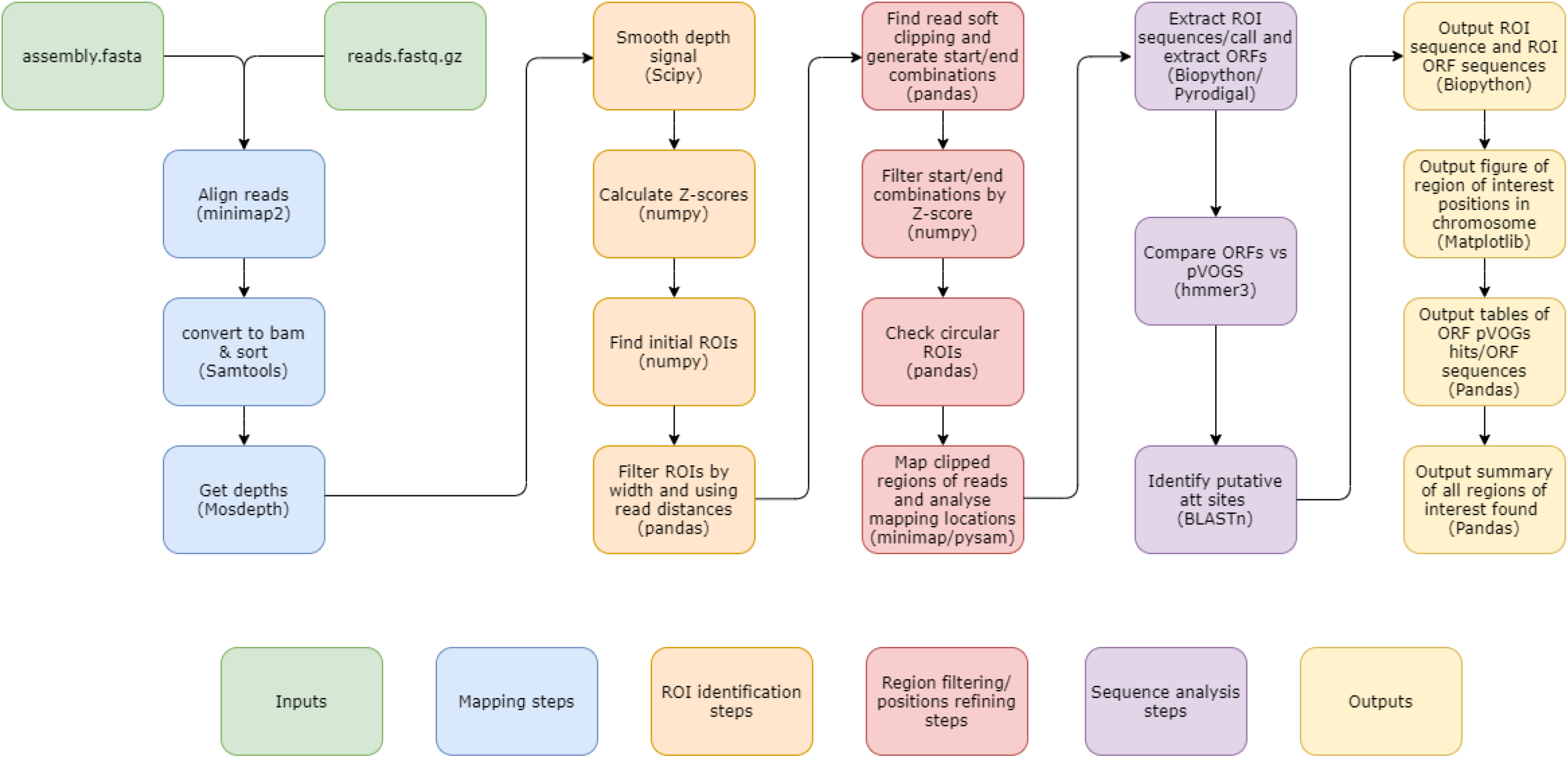
Overview of steps involved in the hafeZ pipeline.

**Figure S2:**
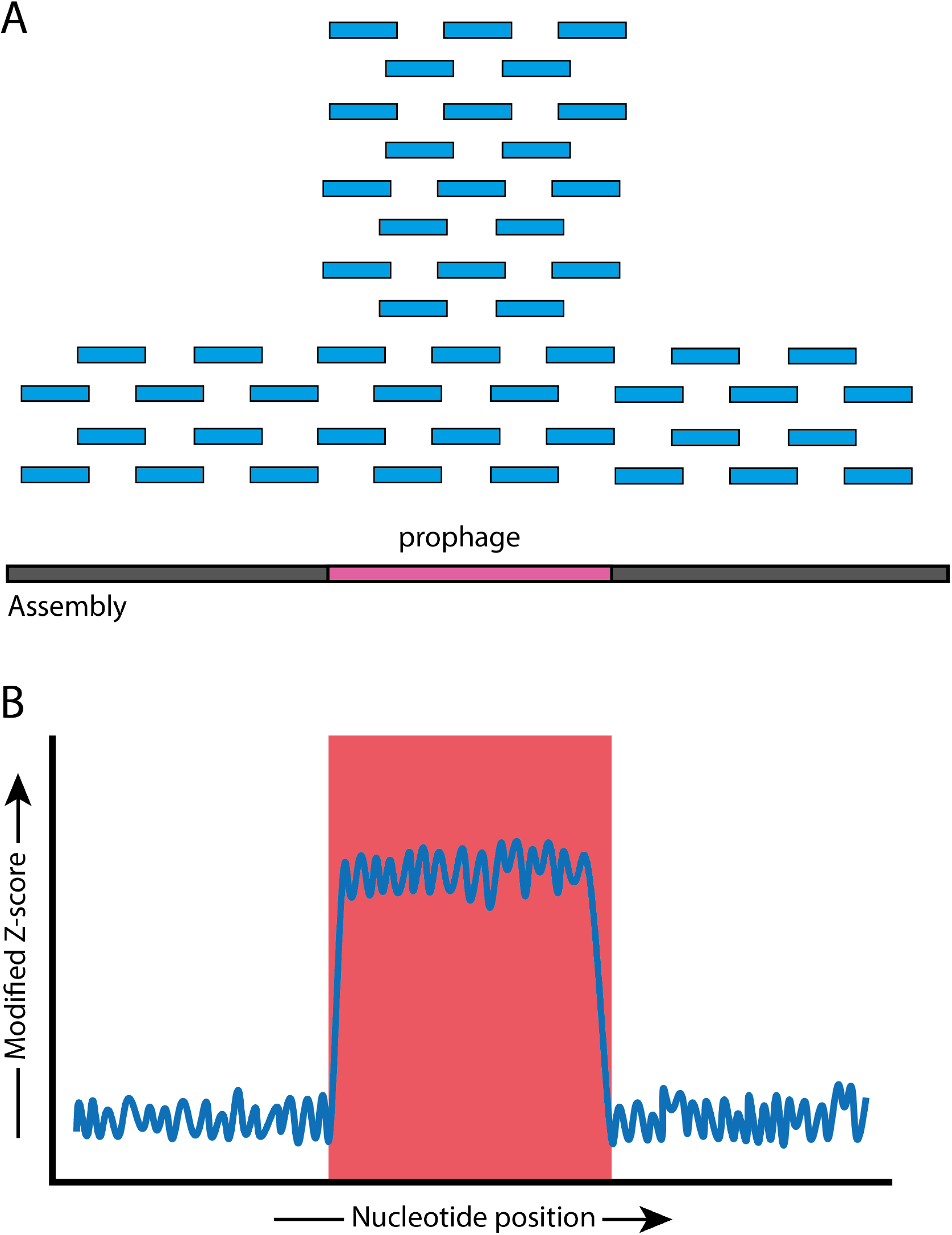
Illustration of how active prophages can be detected from read mapping coverages. A. When reads (blue bars) are mapped to an assembly (grey bar), if a prophage (pink region) exists within the genome that has been induced, a higher frequency of reads mapping to the region of the genome containing the prophage would be expected as copies of the viral DNA should be being produced. B. By converting the per base coverage of the read mapping results to modified Z-score values we can detect regions with higher levels of coverage than the surrounding chromosome (red highlighted region) that may indicate prophage induction.

**Figure S3:**
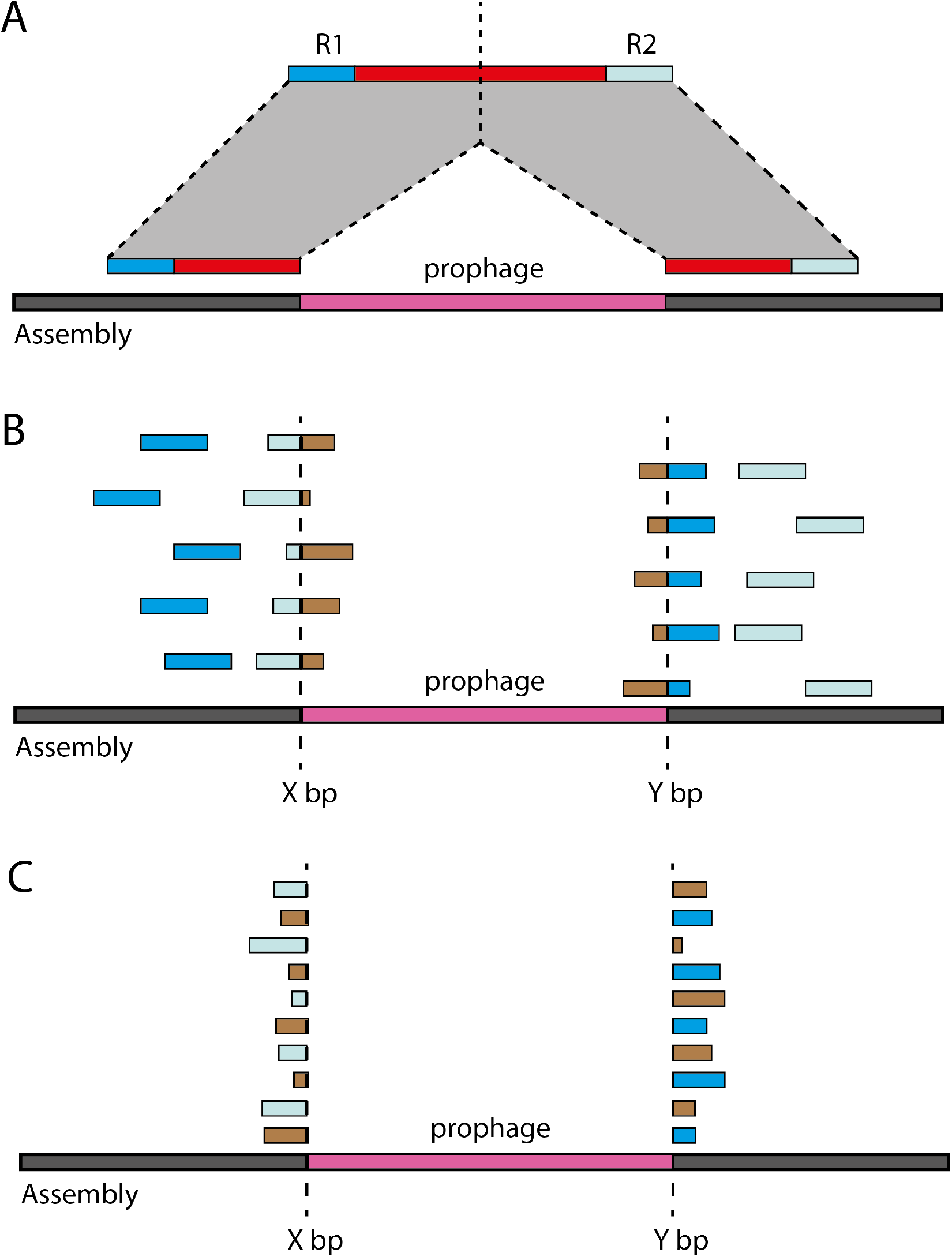
Illustration of how read mapping is used to identify putative active prophages by hafeZ following ROI identification. (A) hafeZ examines reads mapping within an ROI and those mapping within 15,000 bp either side of it for distantly mapping reads. Such reads would indicate that the DNA fragment (red bar) sequenced by a pair of reads (blue ends) contains a deletion compared to the DNA sequence of the genome assembly (dark grey bar) being mapped to. (B) hafeZ inspects the reads mapping within an ROI for the presence of soft-clipped reads, reads where only part of a read maps to the assembly at a given location (blue regions at dotted line = mapped region, brown = clipped region). If a sufficient number of these occur at a consistent bp position (dotted line) this position is then taken as either the start/end of the ROI. (C) By default hafeZ then remaps the regions soft clipped by the initial read mapping step to determine if these reads map at the opposing end of the ROI. This would indicate that the read has sequenced the deleted region where the prophage has excised from.

**Figure S4:**
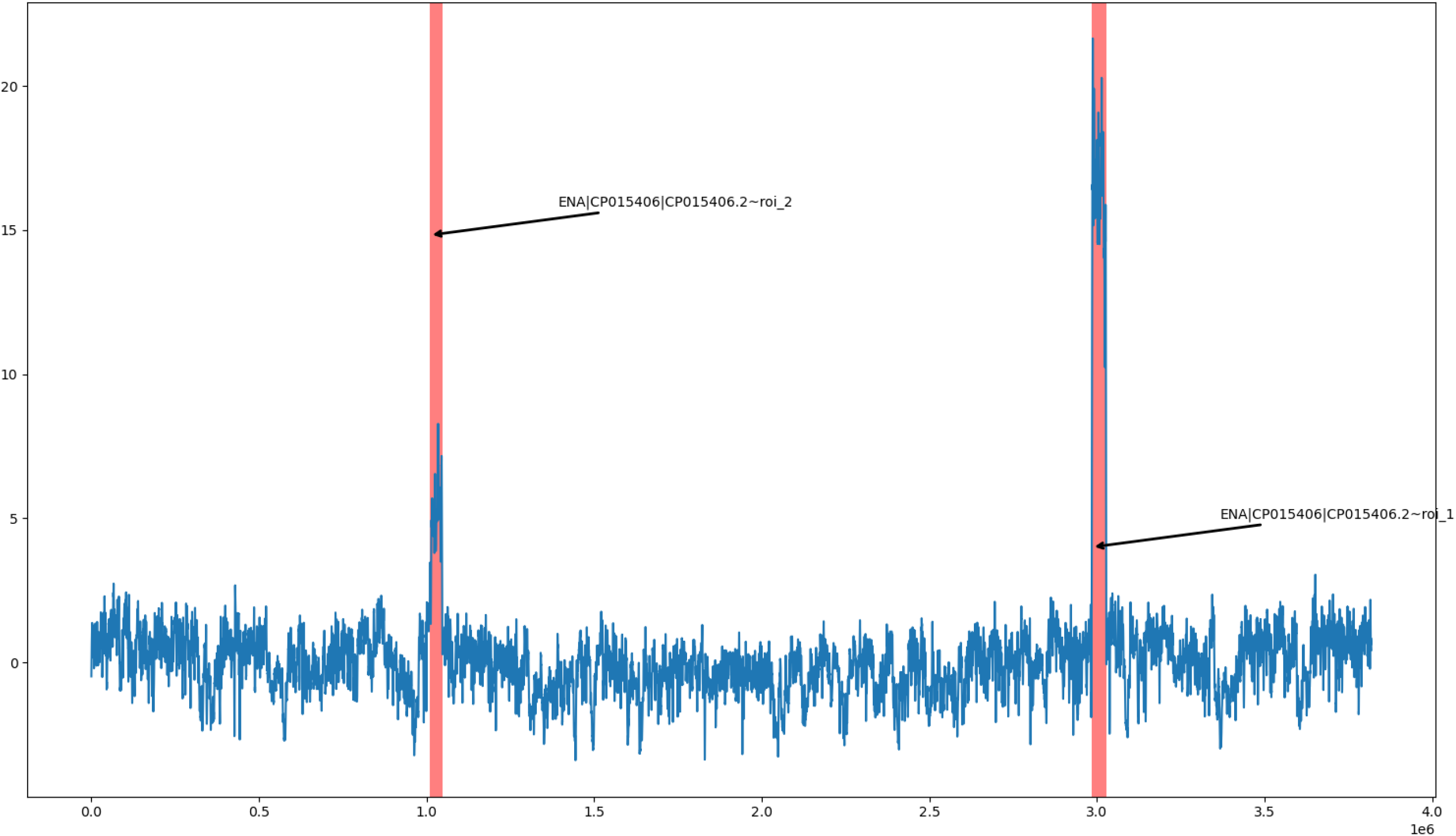
Example of the ROI position figure output by hafeZ for sample YL31_C1. Figure shows modified Z-score on Y-axis and base pair position on X-axis. Each red area signifies the position of a putative active prophage.

Table S1: List and pairings of genome assemblies and reads examined for each sample in this study.

Table S2: Comparisons of the expected positions for the 10 prophages identified by Zünd et al. 2021 and the hafeZ identifications for each sample. Note - The assembly used by Zünd et al., for YL58 could not be found in public databases. Therefore, although the table shows that a new prophage was identified and the expected prophage in YL58 was not identified, we used BLAST to compare the Zünd prophage sequence to the assembly used here and found that the Zünd prophage mapped to the exact start and end positions identified by hafeZ.

Table S3: Table containing the combined output of all summary tables produced for each sample’s hafeZ run.

